# Shared genomic variants: identification of transmission routes using pathogen deep sequence data

**DOI:** 10.1101/032458

**Authors:** Colin J. Worby, Marc Lipsitch, William P. Hanage

## Abstract

Sequencing pathogen samples during a communicable disease outbreak is becoming an increasingly common procedure in epidemiological investigations. Identifying who infected whom sheds considerable light on transmission patterns, high-risk settings and subpopulations, and infection control effectiveness. Genomic data shed new light on transmission dynamics, and can be used to identify clusters of individuals likely to be linked by direct transmission. However, identification of individual routes of infection via single genome samples typically remains uncertain. Here, we investigate the potential of deep sequence data to provide greater resolution on transmission routes, via the identification of shared genomic variants. We assess several easily implemented methods to identify transmission routes using both shared variants and genetic distance, demonstrating that shared variants can provide considerable additional information in most scenarios. While shared variant approaches identify relatively few links in the presence of a small transmission bottleneck, these links are highly confident. Furthermore, we proposed hybrid approach additionally incorporating phylogenetic distance to provide greater resolution. We apply our methods to data collected during the 2014 Ebola outbreak, identifying several likely routes of transmission. Our study highlights the power of pathogen deep sequence data as a component of outbreak investigation and epidemiological analyses.

## Introduction

Genomic data offer new insights into epidemiological and evolutionary dynamics, and sequencing pathogen samples is becoming increasingly routine. Pathogen genomic data allow us to determine the phylogeny of isolates, which in turn sheds light on the potential transmission networks between the hosts from whom they were collected. As such, inference of transmission trees using genomic data is an increasingly well-studied field (1-7). While low-resolution pathogen typing has been used for some time to discriminate between independent outbreaks (8-10), whole genome sequencing provides additional resolution with which genetic distance between identical phenotypes may be ascertained (11-13). This too, however, has limits. Studies have shown that while transmission clusters may be identified with genomic data, individual-level transmission routes can rarely be identified with a great degree of certainty (2, 3). Characterizing an infected host by a single pathogen genome (isolation and purification of a single colony for bacteria, or using the consensus sequence for viral pathogens) is common practice, yet neglects within-host diversity. The variation in sampled genetic distances can be large relative to the expected number of mutations between hosts, rendering the number of SNPs a rather crude measure of relatedness on an individual level (14). As such, particularly for rapidly evolving pathogens, or those whose mode of transmission is associated with a large and potentially diverse inoculum (‘transmission bottleneck’), single genome sampling can cause hosts to appear misleadingly similar or dissimilar.

Deep sequencing can potentially provide new insights into within-host diversity. Currently, sequencing a mixed population sample to sufficient depth to identify minor nucleotide variants has mostly been limited to viral samples. While consensus sequences may appear identical for two samples, comparing minor variants can offer additional resolution. For instance, if the same nucleotide variation is observed at the same locus in pathogen samples from two individuals (henceforth referred to as a ‘shared variant’ (SV)), this could be considered as strong evidence for direct transmission, particularly if the variant is not observed in any other host. This naturally relies on the possibility that a pathogen population of size greater than one survives the transmission bottleneck; otherwise each infection must initially be monoclonal, implying that any variation found within distinct hosts must have arisen independently. There is evidence for larger bottlenecks occurring in viral pathogens such as influenza (15, 16) and Ebola (17), and it is plausible that this is also the case for bacterial pathogens (18).

The connection between SV presence and direct transmission has previously been suggested. Gire et al. noted the presence of SVs in Ebola virus samples from individuals who were potentially linked by transmission (19). Data collected from two influenza A animal transmission studies were used to explore the presence of SVs between hosts, and it was shown that such data were consistent with known contact patterns (20). This study used known contact patterns to identify characteristics of SVs which were more likely to be associated with transmission, allowing variants to be split into those consistent and inconsistent with transmission, minimizing false connections. Poon et al. identified routes of influenza transmission occurring during a household contact study using both consensus whole genome sequence data and the presence of SVs (21). In the case of bacterial pathogens the diversity in *S. aureus* infections, which can be considerable, has been linked to transmission in a veterinary hospital (22).

Pathogens vary considerably in their bottleneck size, mutation rates and transmission dynamics. It remains unclear how methods based on SVs are expected to perform in different regions of this parameter space. Establishing this is a crucial component of the interpretation of SVs and the value of the approach.

In this study, we investigated the predictive power of SVs for identifying transmission routes. In addition to pathogen genomes, many other sources of data many contribute information towards inference of transmission routes, including temporal and spatial data, contact patterns and expression of symptoms. However, here we aimed to examine the information contributed by genomic data alone and in particular the additional benefit offered by considering SVs.

### Methods

We generated infectious disease outbreaks with within-host pathogen evolution under various mutation rates and bottleneck sizes by simulation. We expanded upon methods previously used to infer transmission routes using deep sequence data (19-21), comparing their performance with analogous genetic distance based approaches. We additionally proposed hybrid approaches which combine SV information with phylogenetic distance data. We considered the following approaches:

a. *Weighted variant tree*. For each host, we weight potential sources by the number of observed SVs, such that the host sharing the largest number of variants is attributed the largest weight. Hosts sharing no variants with any other are not assigned a source. Weighting edges provides an extension to previous approaches (19-21).
b. *Maximum variant tree*. For each host, we define the source to be the individual with whom the largest number of SVs are observed. Hosts sharing no variants with any other host are not assigned a source.
c. *Weighted distance tree*. Using consensus sequences, the genetic distance (number of single nucleotide polymorphisms [SNPs]) between isolates is calculated, and potential sources are weighted inversely by this metric. This approach has been described previously (2).
d. *Minimum distance tree*. Using consensus sequences, the source of a given host is defined to be carrier of the genetically closest isolate to that of the host. This approach, with the incorporation of sampling times to provide directionality, has been described previously (6).
e. *Hybrid weighted tree*. First, the weighted variant tree is constructed. Hosts with no source are then assigned potential sources based upon weighted genetic distance.
f. *Hybrid maximum tree*. First, the maximum variant tree is constructed. Hosts with no source are then assigned potential sources based upon minimum genetic distance.

These six simple heuristics by no means comprise an exhaustive list of approaches to identify routes of transmission, but are instead a range of readily-implemented distance-based approaches, which require neither knowledge of evolutionary dynamics, nor infection or sampling times. As has previously been demonstrated, simple methods based on genomic data alone can provide powerful insights into transmission dynamics (23). Further details of the approaches used here, as well as the metrics used to assess the accuracy of tree reconstruction and reliability of estimated transmission routes, are provided in the Appendix. We additionally applied SV methods to previously published data collected during the Ebola virus outbreak in West Africa in 2014 (19, 24).

### Results

#### Simulation Studies

As expected, the proportion of cases in which a SV was observed in at least one other host increased rapidly with mutation rate and bottleneck size (Figure 1A). The majority of SVs were observed in exactly two individuals, with the proportion shared among larger groups declining rapidly as the size of the group increased (Figure 1B). For each simulation, we constructed a weighted transmission tree according to the six methods outlined previously. An example simulated outbreak of a pathogen with similar characteristics to *S. aureus* (see Appendix) is shown in Figure 2, along with reconstructions based upon two of these methods. While many edges are bidirectional and symmetric, asymmetry can occur under most methods due to the lack of commutativity – i.e. even if B is the closest host to A, A may not be the closest host to B.

**Figure 1.**
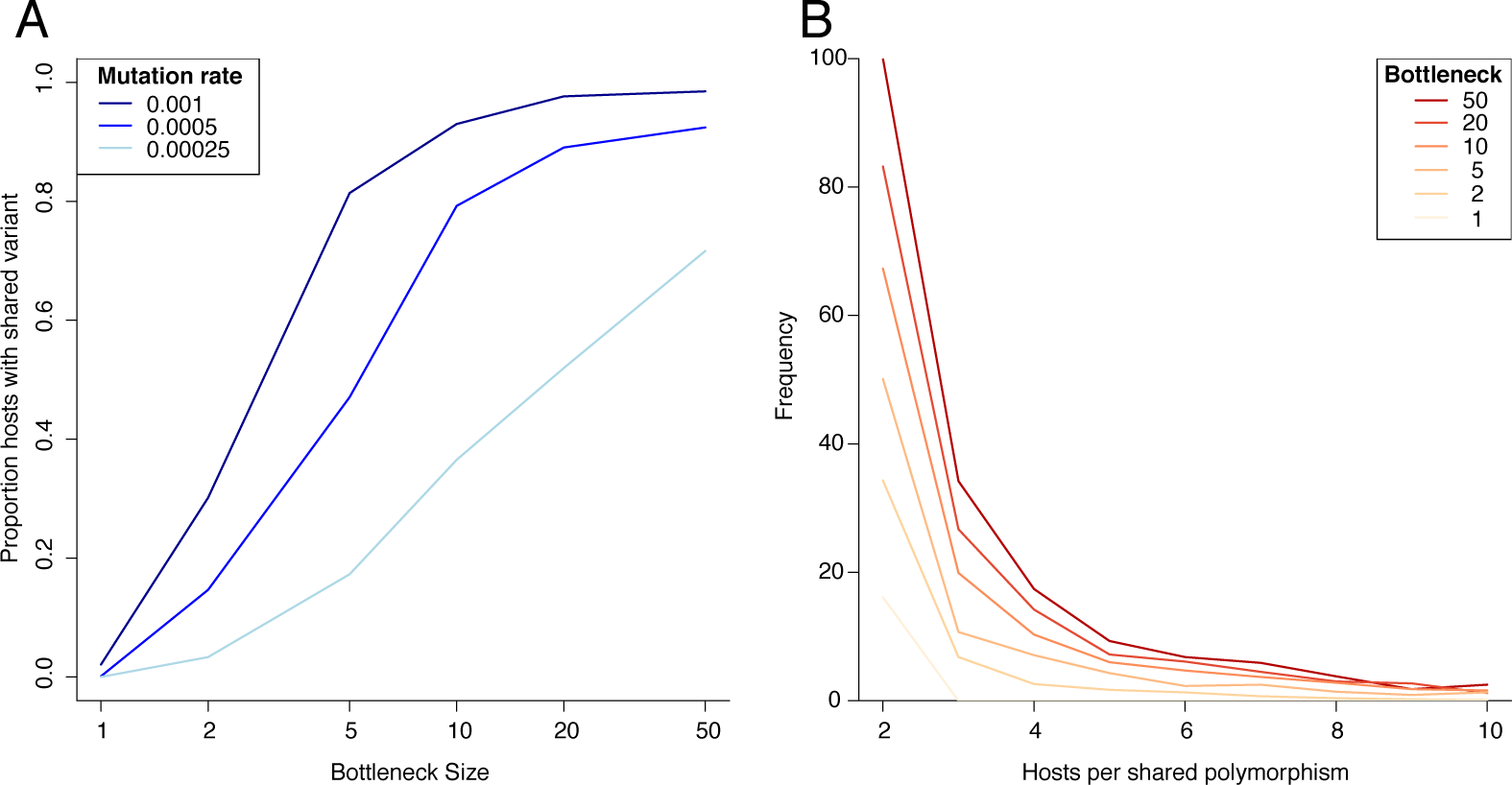
Summary of genetic variant frequency across the simulated outbreaks. We simulated 10 outbreaks for each combination of six bottleneck sizes and three mutation rates (180 in total). (A) Mean proportion of cases in outbreak for whom at least one shared variant was observed. (B) Distribution of shared variant group size for different bottlenecks, with a mutation rate of 5×10^−4^ per genome per generation.

**Figure 2.**
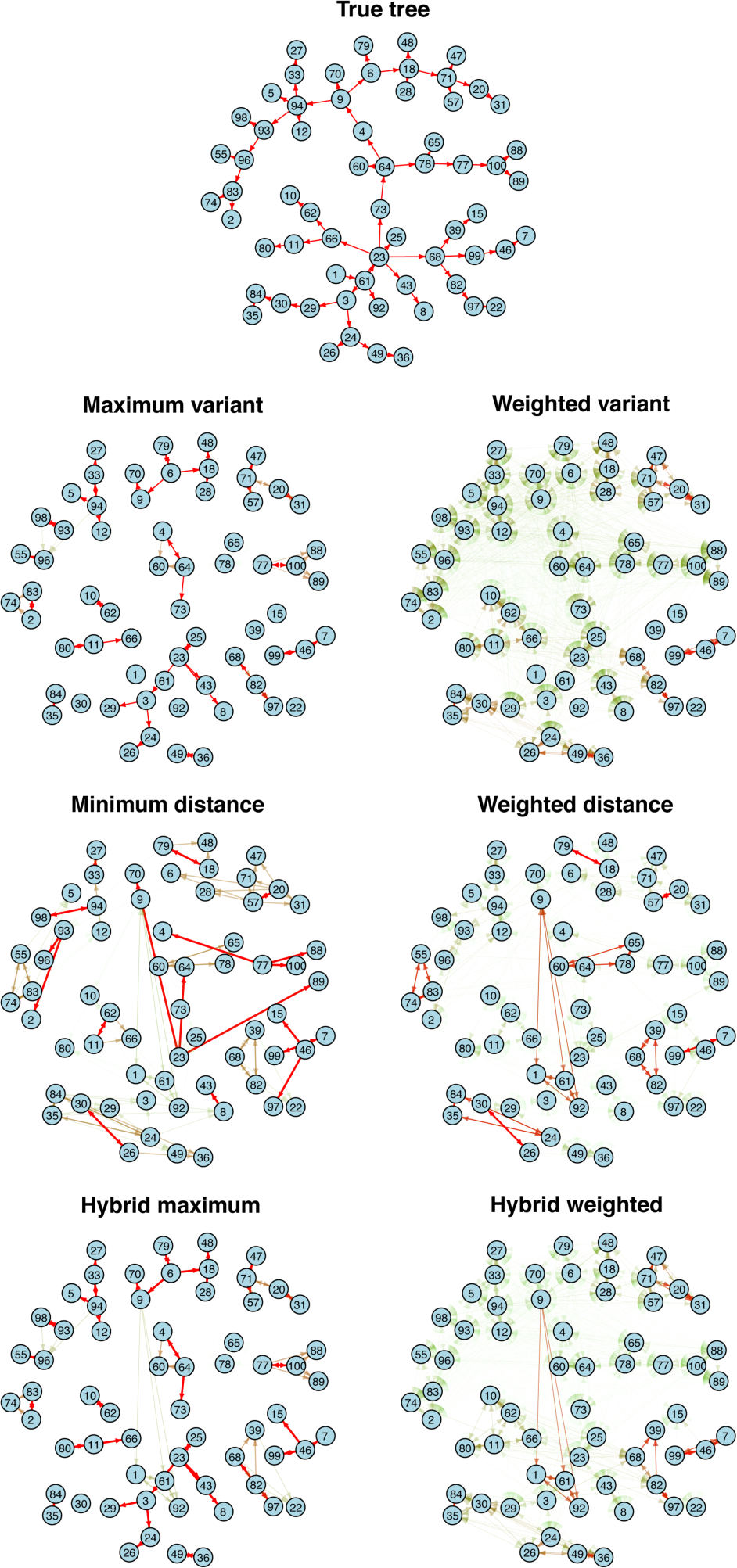
Simulated and reconstructed transmission trees. The simulated tree (A) was generated with bottleneck size 10 and mutation rate 0.001 per genome per generation. Trees were reconstructed according to the maximum variant (B) and the minimum genetic distance (C) approaches described in *Methods*. Edges are weighted according to the weight attributed to that potential transmission route. Networks were plotted with the igraph package in R.

We used two metrics to assess the reliability of individual estimated transmission routes and compare the different methods described above. Firstly, we considered the true path distance between inferred transmission pairs. We found that under the maximum variant tree, the mean path distance was typically less than 2, outperforming the minimum distance approach (Figure 3A). Secondly, we examined the mean weight assigned to the true source of each host. In the case of small (<5) bottleneck sizes, methods based on SVs perform poorly since the likelihood of a monoclonal infection is high, resulting in most true links being assigned a weight of zero. Furthermore, those links which are inferred by SVs in small-bottleneck settings identify direct transmission with high confidence. The hybrid approaches perform best for small bottlenecks, incorporating SV information when available but not relying upon it. For larger bottlenecks, the distance-based approaches were markedly outperformed by the variant-based approaches (Figure 3B). As the rate of mutation increases, SV approaches outperform those based on distance alone to an increasing degree as mutation generates increasing diversity in the infecting population (Figures S1A-C).

**Figure 3.**
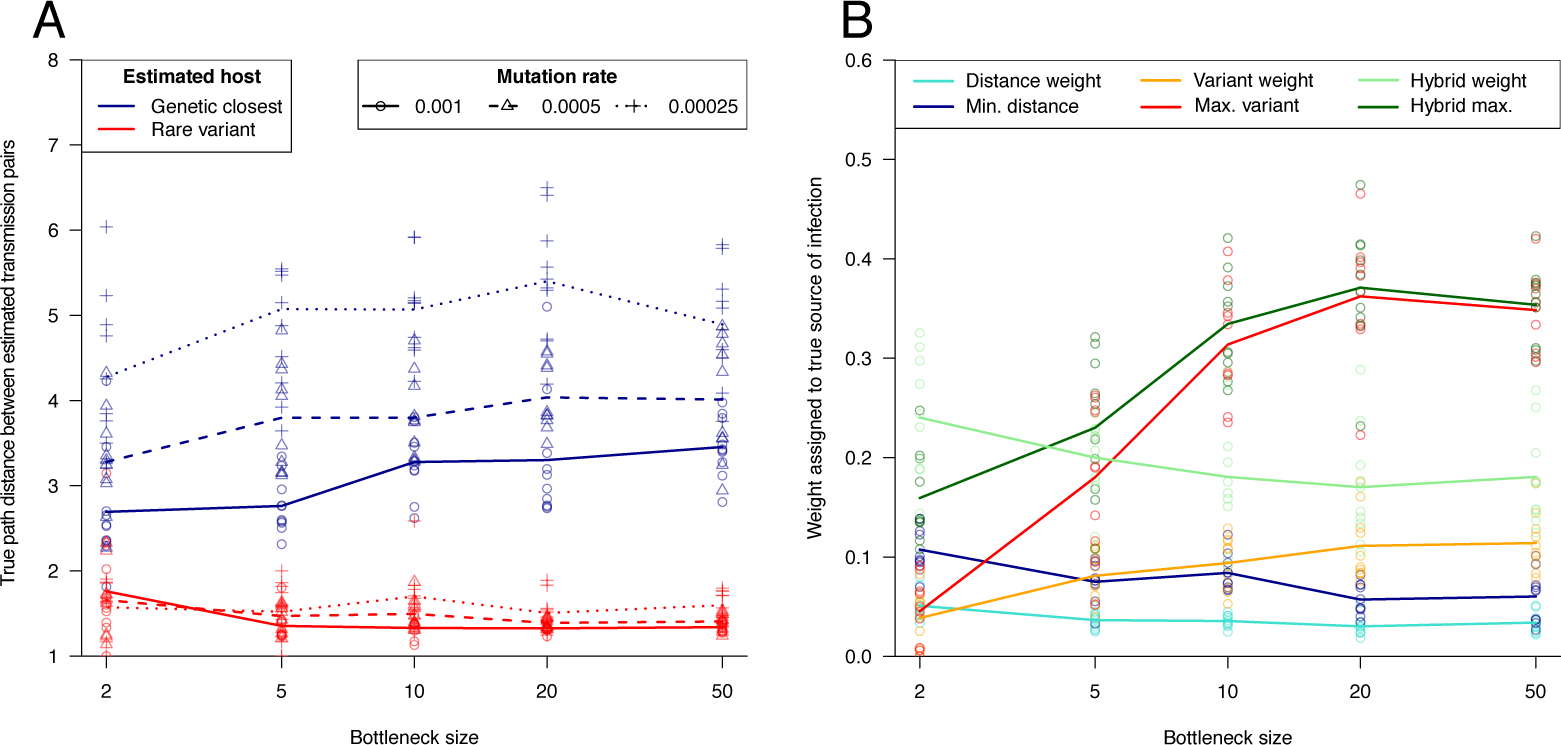
Reliability of estimated transmission routes. (A) The true path distance between estimated transmission pairs gives insight into the extent to which transmission links are misspecified. Here we compare the minimum distance (black) and the maximum variant (gray) approaches. A perfect reconstruction would have mean path length 1. Maximum variant path lengths are averaged over identified transmission pairs, that is, excluding hosts with no shared variants. (B) The mean weight attributed to each true transmission link for each tree reconstruction method, under a range of scenarios and methodologies. Results are shown for a mutation rate of 5×10^−4^ per genome per generation, and were averaged across 10 simulated outbreaks for each scenario.

In addition to the reliability of individual links, we also considered the overall accuracy of a transmission tree reconstruction. This was measured by the area under the receiver operating characteristic curve (AUC) statistic. For small bottlenecks, variant-based methods provide a poor tree reconstruction by this metric (Figure 4); values below 0.5 indicate a worse performance than random selection; an inevitability when only a small proportion of nodes are assigned sources. A tight bottleneck leads to little diversity persisting across transmission events, and as such, SVs are rarely observed, leading to a sparsity of informed links across the network. However, larger bottlenecks lead to rapidly improving AUC statistics for the variant-based approaches, which even exceed the weighted distance approach with a sufficiently large bottleneck size and mutation rate (Figure 4, Figures S1D-F). In contrast, distance-based approaches typically decline in accuracy as the bottleneck size increases, for reasons that are well understood (2, 25) We additionally investigated the effect of ‘mutational hotspots’, which can generate potentially confounding homoplasy. We found that while variant approaches performed less well, they generally continued to outperform distance based approaches for larger (>10) bottleneck sizes (see Appendix, Figures S2 & S3 for further details).

**Figure 4.**
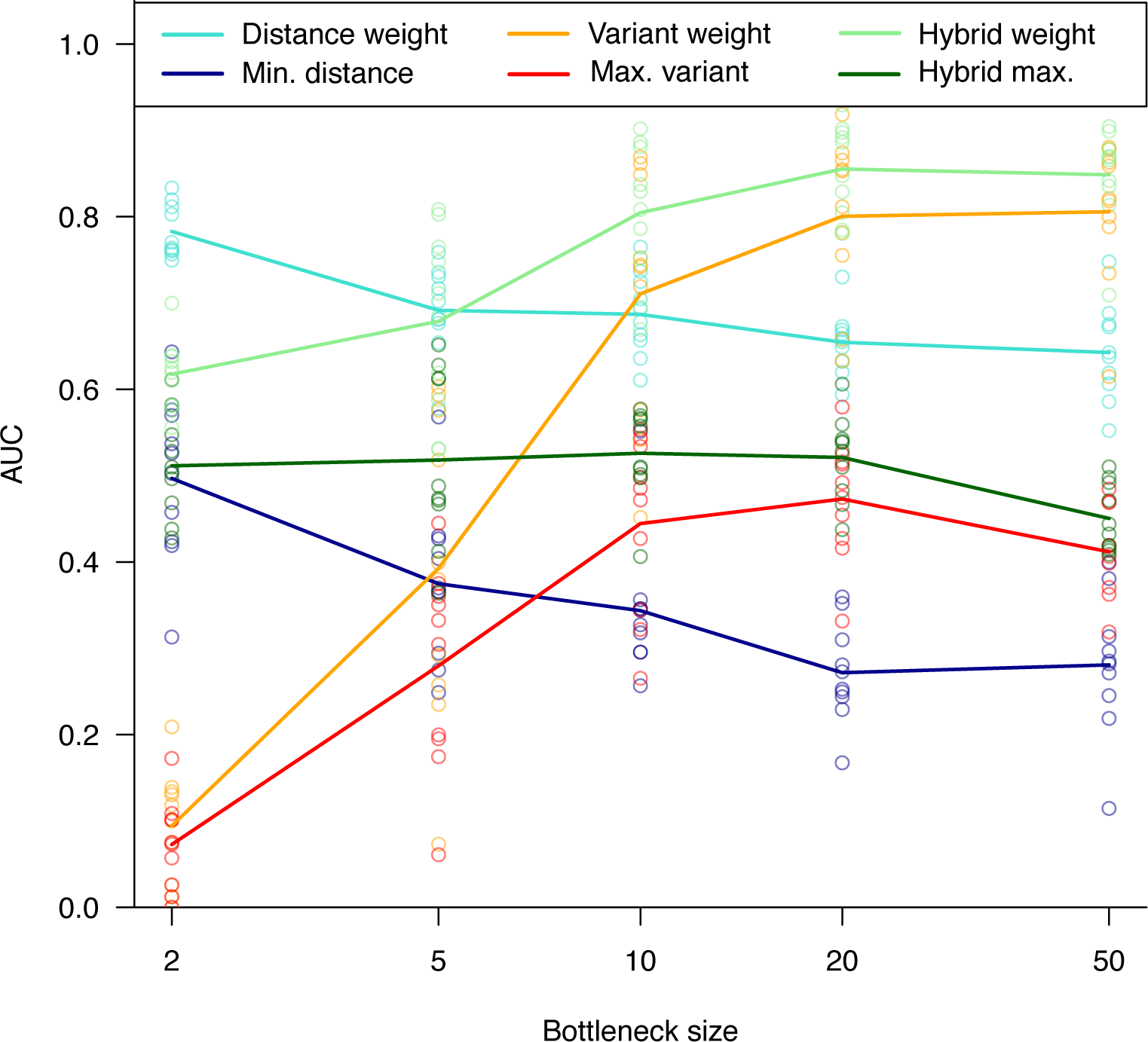
Transmission tree reconstruction accuracy. The area under the receiver operating characteristic curve (AUC) metric provides an overall measure of network accuracy. Results for a mutation rate of 5×10^−4^ per genome per generation are shown here. The mean AUC across 10 simulations was measured for each scenario.

#### Ebola virus data

We next examined previously published Sierra Leone Ebola datasets, for which raw reads are available and we can determine the presence and properties of intra-host variants. In order to reduce the risk of counting variant calling errors as true intra-host variants, we identified only variants in which the minor frequency was at least 5% (routes estimated under a 1% threshold are shown in Figure S4). Figure 5 shows the transmission trees reconstructed for each dataset under the weighted variant approach, using no epidemiological information. In the first dataset (Figure 5A, (24)) 19/78 hosts were found to share a variant with at least one other individual. Four pairs of patients shared more than one variant (three pairs with two SVs, and one pair with four), while one additional pair shared one unique variant. Consistent with transmission, each of these pairs originated from the same geographic location, and permutation testing revealed this geographic similarity was significantly higher than would be expected via random selection (*P*=0.0075). Pairs were also temporally clustered; three of these links were sampled two or fewer days apart, while the remaining two were sampled 12 and 22 days apart, which are plausible given serial interval estimates for Ebola virus infection of 15.3 ± 9.3 days (26). Under the minimum distance tree, two of these pairs were reproduced, two pairs belonged to a much larger group of samples with identical consensus sequences, and one pair, differing by one SNP according to consensus sequences, remained unconnected due to the presence of other identical sequences (Figure S5).

**Figure 5.**
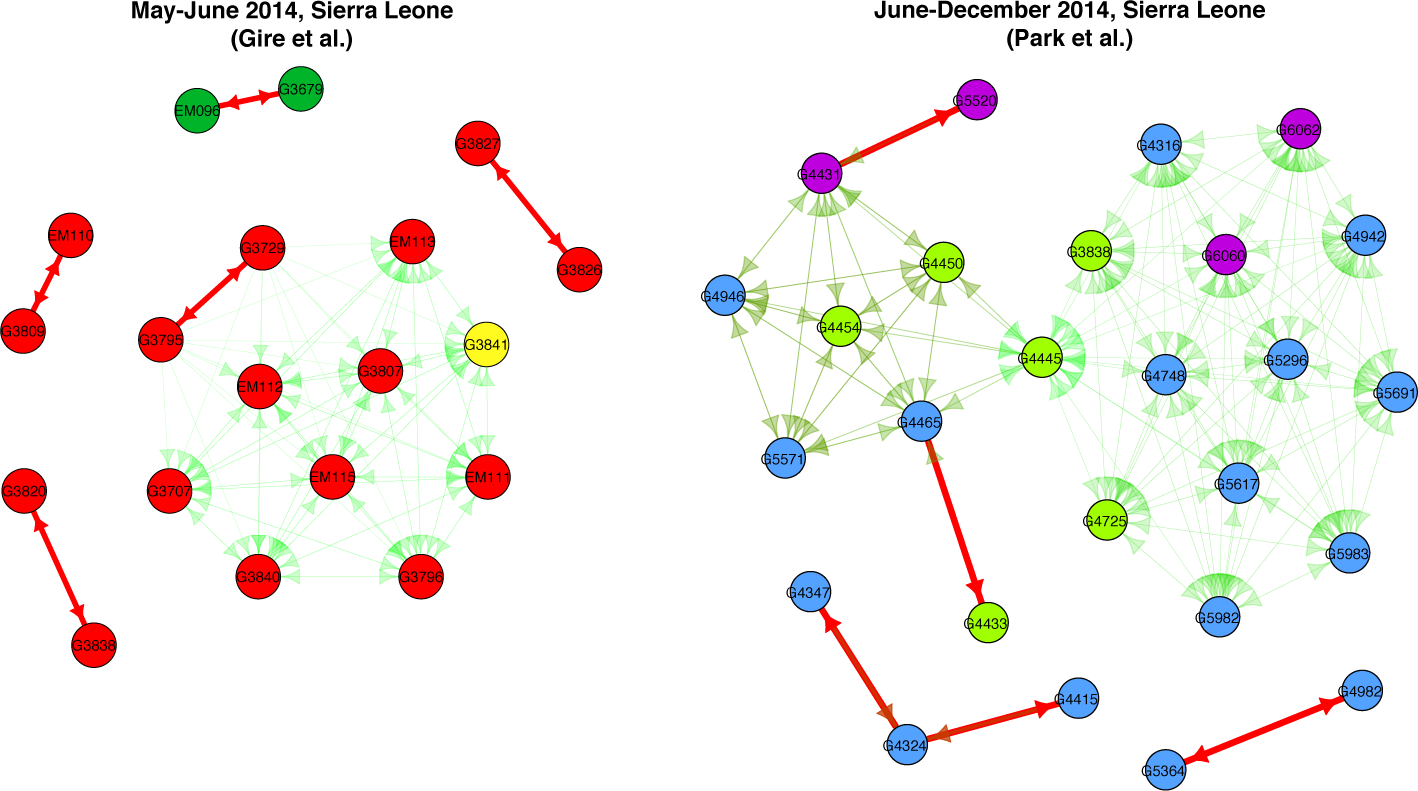
Estimated Ebola transmission routes. Transmission links between sampled hosts in the 2014 Ebola outbreak under the maximum variant approach. Colors denote chiefdoms to which hosts belong, while the weight and thickness of the arrows denote the relative weight attributed to each potential transmission event, with bold arrows denoting a higher weight than light arrows. Variant detection threshold 5%.

While a consistent result of our simulations was the sharing of variants among small numbers of hosts, rarely more than two, in the Ebola data collected by Gire et al., one variant was shared by 11 hosts. Samples with this variant are highly clustered geographically (10/11 same chiefdom, *P*=0.022) and temporally (observed within 18 day period), as well as phylogenetically (19), lending support to this group representing a transmission cluster.

In the second dataset (Park et al.) 26/150 (for which replicate sequencing and variant calling was performed) shared a variant with at least one other host (Figure 5B, (24)). There were five pairs of individuals sharing a unique variant. As before, one variant was shared by multiple hosts, but unlike the previous dataset these were not geographically or temporally clustered, coming from different villages and spanning several weeks. Furthermore, while some of these samples cluster on the phylogenetic tree, many fall in different clades (24).suggesting the group is unlikely to represent a single transmission cluster, rather multiple transmission events in combination with homoplasy. Four of the five ‘unambiguous’ transmission pairs joined patients from the same geographic location, consistent with transmission, significantly greater similarity than would be expected by random selection (*P*=0.0073).

### Discussion

We have described some simple methods for reconstructing transmission trees using shared variants, testing how well this approach performs for a range of parameters governing the rates of diversification within and between hosts. We have then applied the methods to data from the recent Ebola outbreak to ask whether it identifies links, using the genomic data alone, that are likely to be consistent with transmission given time and location.

For the great majority of parameter space, excluding only very low mutation rates and tight bottlenecks, these methods outperform genetic distance comparison methods, which have increasingly been used to identify potential transmission events (6, 27, 28). The limitations of distance-based methods that characterize a single genome are well appreciated. We note that while for the purpose of comparison, additional data sources were not included in our inference of transmission routes, incorporating these independently would be a relatively straightforward step with these methods. Most simply, sampling dates could be used to provide directionality to inferred connections.

The additional information we derive from SVs can inform the transmission tree in two distinct ways, depending on the region of parameter space. Firstly, small but non-singular bottlenecks (eg. for airborne influenza transmission (29), sexually transmitted HIV (30)) produce few inferred transmission pairs, but these are highly accurate. The small bottleneck means that the probability of observing a SV between individuals who are in the same transmission cluster, but not directly linked, is negligible. Secondly, SV data for pathogens with larger transmission bottlenecks (eg. Ebola (17), influenza transmitted via contact (29), intravenous drug associated HIV transmission (31)) provide good information on the overall tree structure and transmission clusters, but individual links may be more uncertain. In all cases, higher mutation rates allow for a greater probability of variants emerging in the first place, and this typically results in better inference of transmission routes.

A hybrid approach that combines SVs and the sequence of either an individual sequenced genome or the consensus, offers substantial benefit in the case of small bottleneck sizes (<5), where we predict a method based on SV alone will struggle. Since transmission routes are assessed independently of one another, estimated transmission trees frequently comprise several unconnected nodes or clusters. Such unconnected clusters could be linked to one another if further structure is required, using the weighted distance approach on pooled within-cluster samples. We have here simply used genetic distance, which is predicted to be efficient and reliable under the relatively short time scale of an outbreak, but more sophisticated models of sequence evolution could be applied.

We applied these methods to Ebola data collected from Sierra Leone in 2014. While the first dataset is thought to represent relatively dense coverage of the initial stages of the epidemic in the country, with estimates of around 70% of cases sampled (19, 32), the later dataset comprised a sparser sample. While sparse sampling reduces the number of true links one would expect to find via any method, the reliability of transmission routes identified via SVs remains largely unaffected (Figures S6 and S7). As such, while only relatively few transmission routes were identified in the datasets, this is likely a function of both the proportion of missing data, and the relatively low mutation rate of Ebola virus (19, 33). Confidence in the transmission pairs identified was reinforced by investigating temporal and geographic clustering, which proved to be significant, and while the aim of our study was to assess the accuracy of transmission route identification via genomic data alone, methodology combining spatial and temporal data sources will naturally provide further insight. Identifying even a small proportion of direct transmission pairs can be of great interest in terms of studying pathogen level transmission dynamics and outbreak investigations.

Studying the Ebola data revealed both datasets contained a large group of hosts sharing the same variant, which was rare in all our simulations. The observation can be explained in at least two ways – recurrent mutation (such as might arise through selection) or an anomalously large number of contacts with large bottleneck size (such as might be associated with a funeral based exposure). Park et al. suggest that the large group in the second dataset likely arose through a combination of patient-to-patient transmission and recurrent mutation (24). Subsets of this group do cluster on the phylogenetic tree, which suggests that an alternative hybrid approach, in which large groups are partitioned by genetic background, may prove insightful.

Sample contamination may be an additional source of error. Cross contamination may potentially lead to shared variants observed between unlinked hosts. However, in most settings we do not believe this would present a major concern. In many cases, contamination will not lead to shared variants. If a genotype from sample A contaminates sample B, they will not be linked if the sample A genotype differs from sample B genotypes only at positions invariant in sample A. If no minor variants are contained in the contaminating sample, shared variants will not link this to the contaminated sample. However, it remains important to verify that observed shared variants are consistent with transmission, and to minimize the risk of contamination occurring as much as possible. As before, one can check the genetic background upon which variants are observed. If consensus sequences differ by more than a small degree, then it is likely that homoplasy or contamination may be a confounding issue.

Another deliberate simplification in the present work is the assumption of neutral evolution. While this is plainly faulty over longer time scales, over the relatively short timescale of an outbreak it is a first approximation, and this is supported by real data from outbreaks (19, 24) and even longer periods (34) showing evidence of incomplete purifying selection. Selection may not however have as severe an effect on these methods as we might assume. If a specific variant is maintained through balancing selection it is likely to be found in multiple hosts, and as a result will be less informative as to specific transmission links; if several hosts are connected by the same SV, this will be misleading only if no additional variants are observed. In contrast diversifying selection at antigens, for example, is expected to produce the mutational hotspots we have studied here, which again have little impact. A similar argument can be made that sequencing errors will be less important than expected, because they are likely to be found in just one sample and hence be uninformative as to links to other samples. A more formal approach to this problem would be to test for selection and down weight the identified loci from the analysis.

As yet, there are still few studies in which adequate data have been collected in order to use SVs as a feature to identify transmission routes. Deep and high quality sequencing is required to reliably call minor variants, as well as dense sampling of the outbreak population such that the majority of infection sources are included in the study population. It is likely that such data will become more commonly collected in the near future, for both viral and bacterial pathogens, as the associated sequencing costs fall and the benefits become more evident. This work should motivate research to determine the mutation rates and bottleneck sizes for more pathogens. It is noticeable that bottleneck size in nature, as opposed to minimal infectious dose, has not received the attention is deserves. The importance of this parameter for these methods, as well as other factors like the evolution of virulence (35) should motivate further study.

We have demonstrated the power of deep sequencing data to identify transmission routes with greater resolution than analogous methods using the genome of a single isolate. We have intentionally omitted the incorporation of additional data sources (such as times of sampling, symptom onset, recovery/death, as well as geographic location and contact tracing) in order to evaluate the information provided by the genomic data alone. While this is impressive in itself, incorporating additional data sources will only improve estimates and allow further potential transmission links to be ruled out. Rigorous collection of epidemiological data remains a crucial component of outbreak investigation, and combining this with deep sequencing and SV analysis can provide unprecedented insight into individual-level transmission dynamics.

---

**Figure S1.**
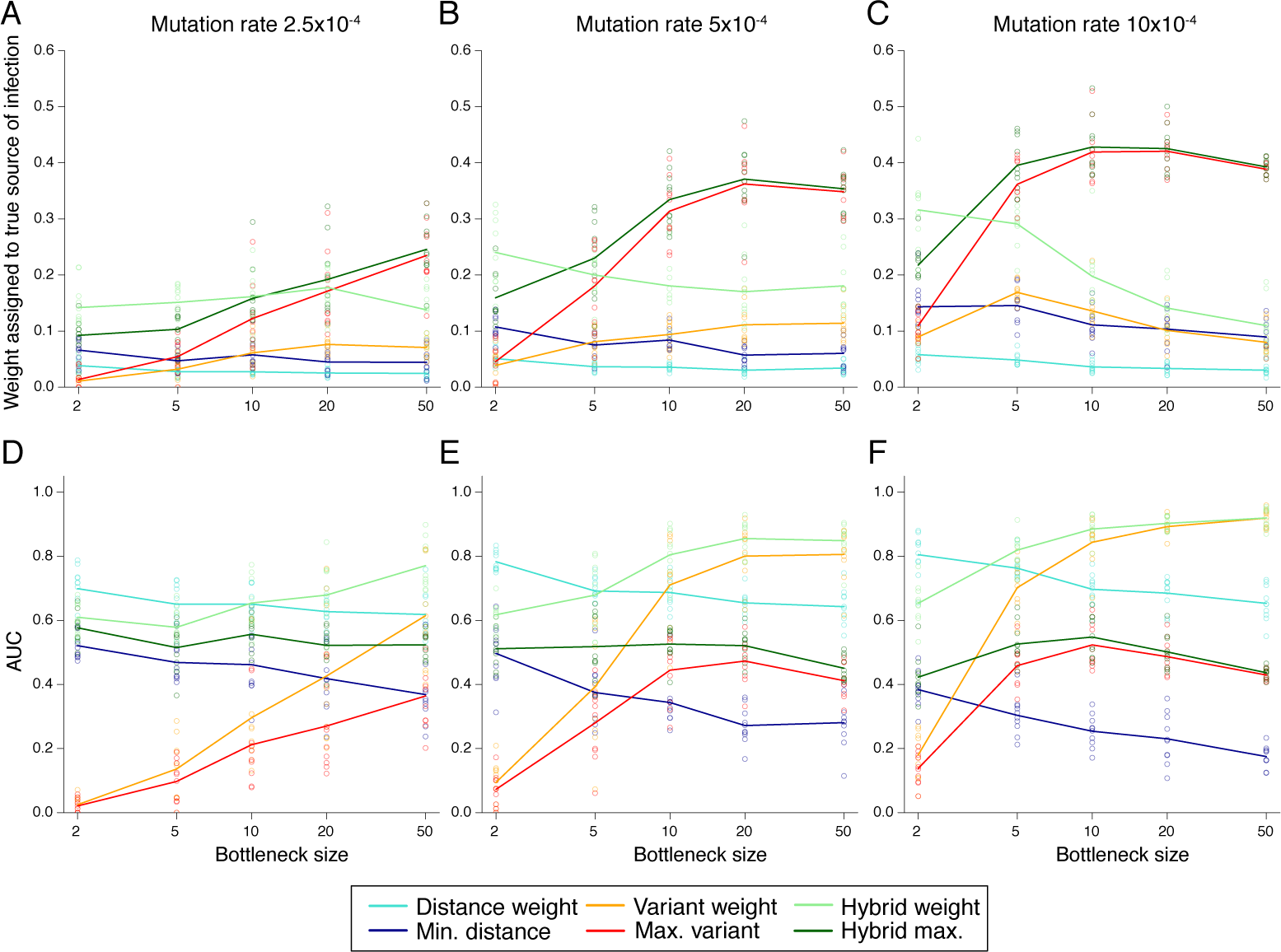
Impact of mutation rates. We generated outbreak data under mutation rates of 2.5, 5 and 10×10^−4^. (A-C) The area under the receiver operating characteristic curve (AUC) metric provides an overall measure of network accuracy. (D-F) The mean weight attributed to each true transmission link for each tree reconstruction method.

**Figure S2.**
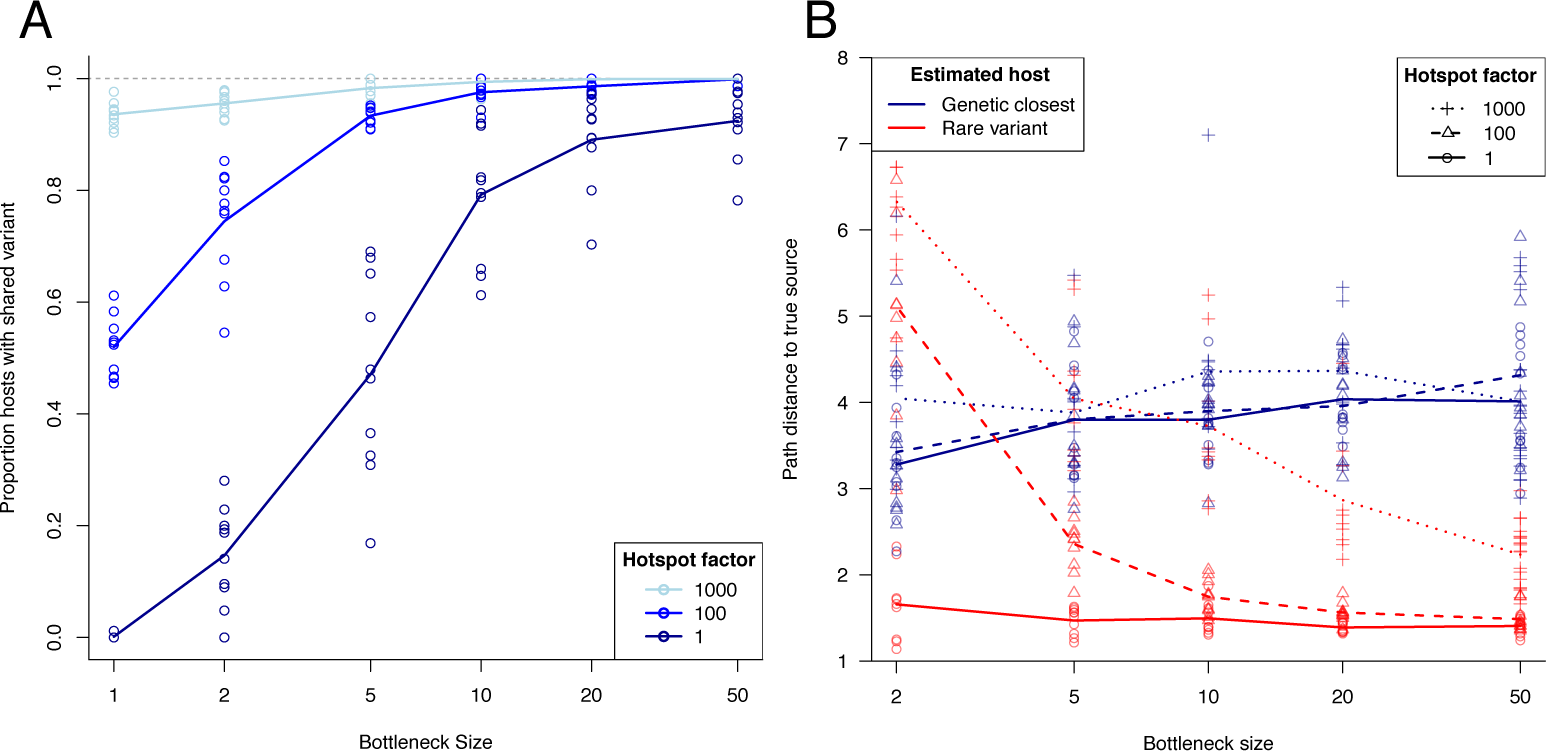
Impact of mutational hotspots. For each scenario, a small proportion of the genome (0.1%) experiences hypermutation, a factor of 1, 100 or 1000 greater than mutation occurs on the remainder of the genome. (A) Proportion of cases in outbreak for whom at least one shared variant was observed. (B) The true path distance between estimated transmission pairs gives insight into the extent to which transmission links are misspecified. A perfect reconstruction would have mean path length 1. Maximum variant path lengths are averaged over identified transmission pairs, that is, excluding hosts with no shared variants.

**Figure S3.**
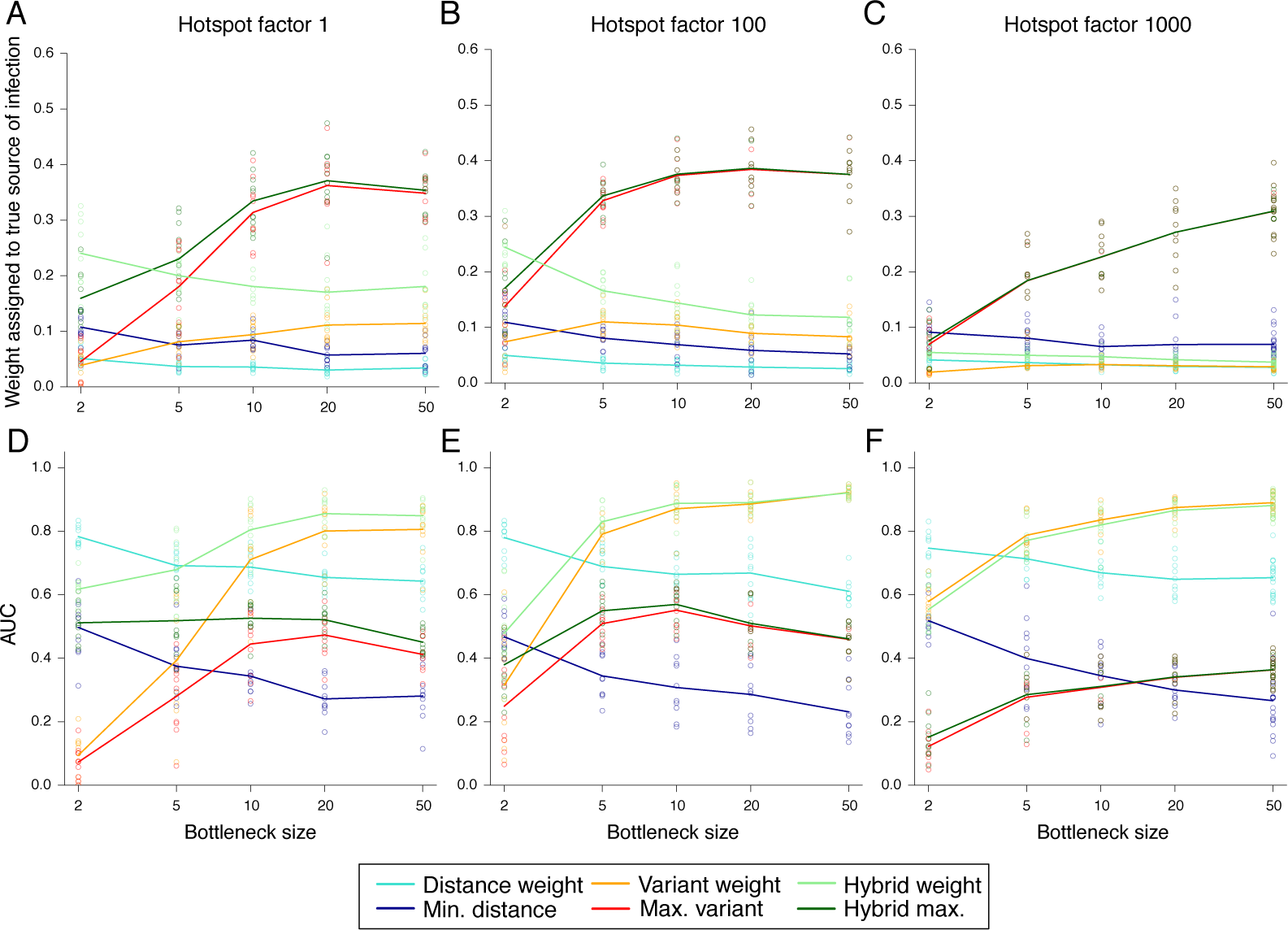
Impact of mutational hotspots. For each scenario, a small proportion of the genome (0.1%) experiences hypermutation, a factor of 1, 100 or 1000 greater than mutation occurs on the remainder of the genome. (A-C) The weight attributed to the true source of infection. (D-F) The area under the receiver-operator curve (AUC), a metric to determine the overall accuracy of network reconstruction.

**Figure S4.**
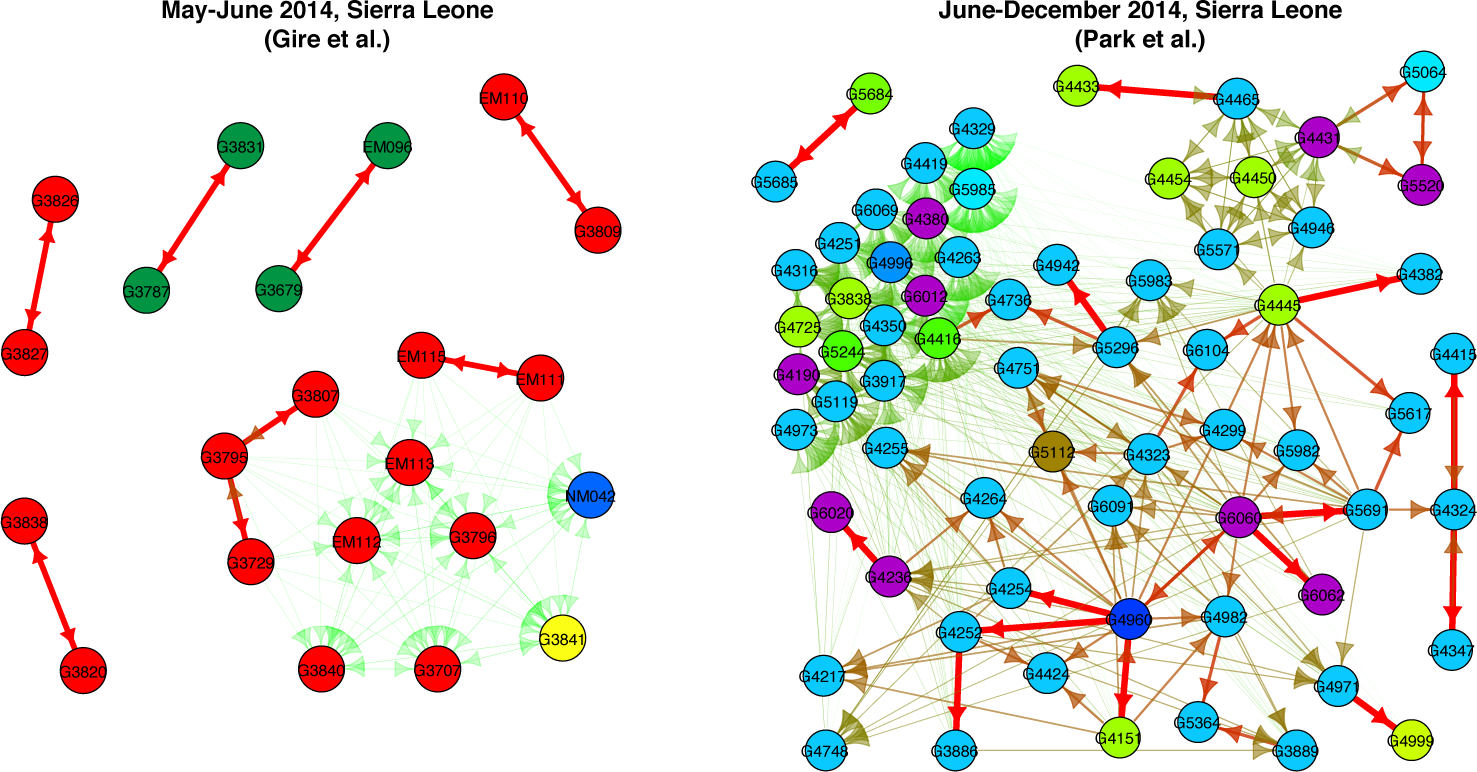
Ebola transmission routes. Estimated transmission links between sampled hosts in the Ebola outbreak under the maximum variant approach. Colors denote the chiefdom to which each host belongs, while the color and thickness of the arrows denote the relative weight attributed to each potential transmission event. Variant detection threshold 1%.

**Figure S5.**
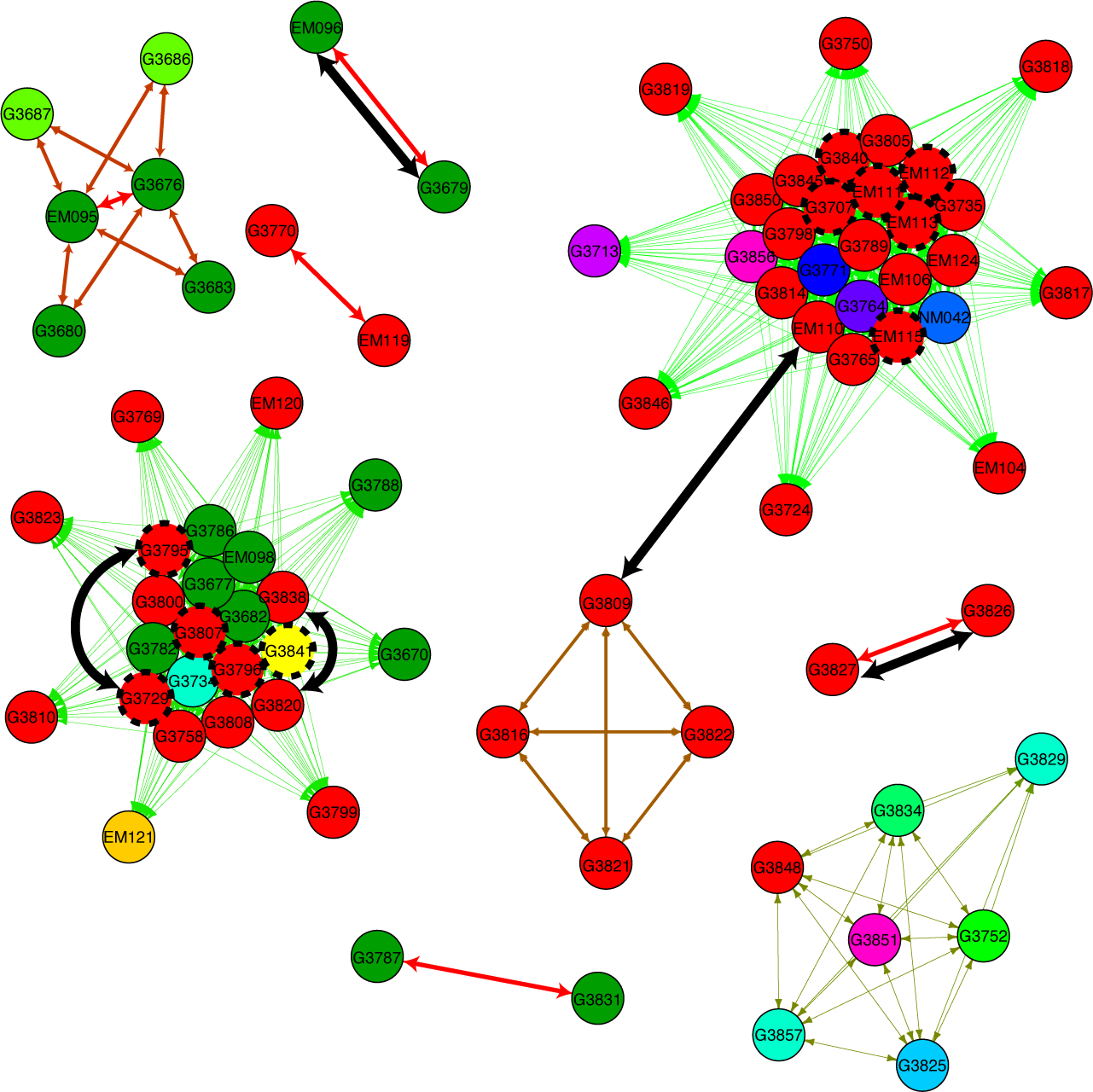
Comparison of Ebola transmission trees using genetic distance and shared variants. All 78 sequenced patients in the first dataset (Gire et al.) are denoted as nodes, colored according to chiefdom. Colored arrows denote edges inferred by the minimum distance approach, with red, bold arrows representing a greater weight than light, green arrows. Black arrows denote edges identified by presence of shared variants, corresponding to Figure 5A. Nodes with a dashed bold outline all share the same variant, corresponding to the large connected group in Figure 5A.

**Figure S6.**
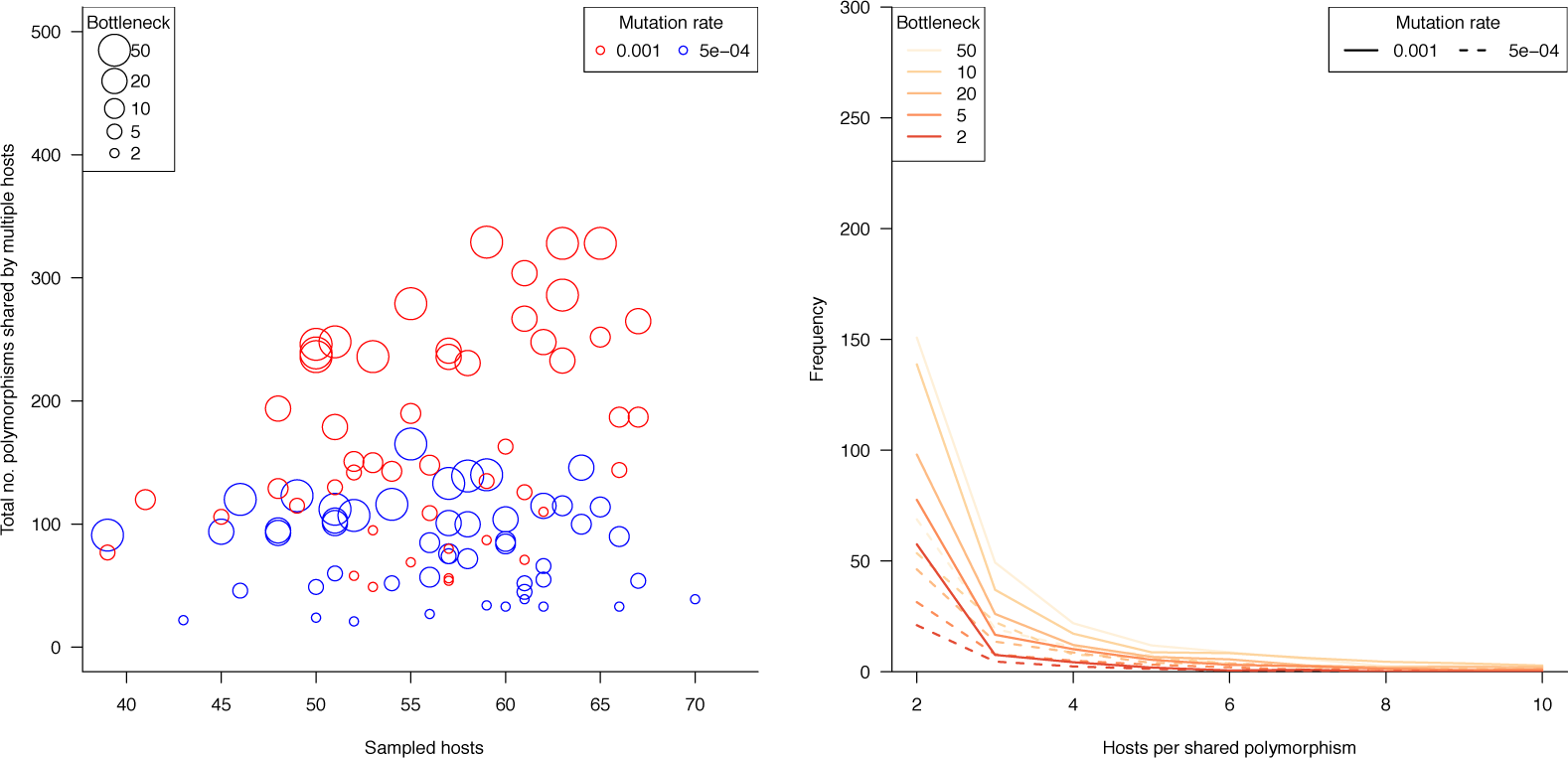
Summary of genetic variant frequency across imperfectly sampled simulated outbreaks. Summary of genetic variant frequency across the simulated outbreaks with 30% of infected hosts unsampled. (A) Total number of shared variants across the simulated outbreak. Bottleneck size is illustrated by circle size. (B) Distribution of shared variant group size for different bottlenecks and mutations rates.

**Figure S7.**
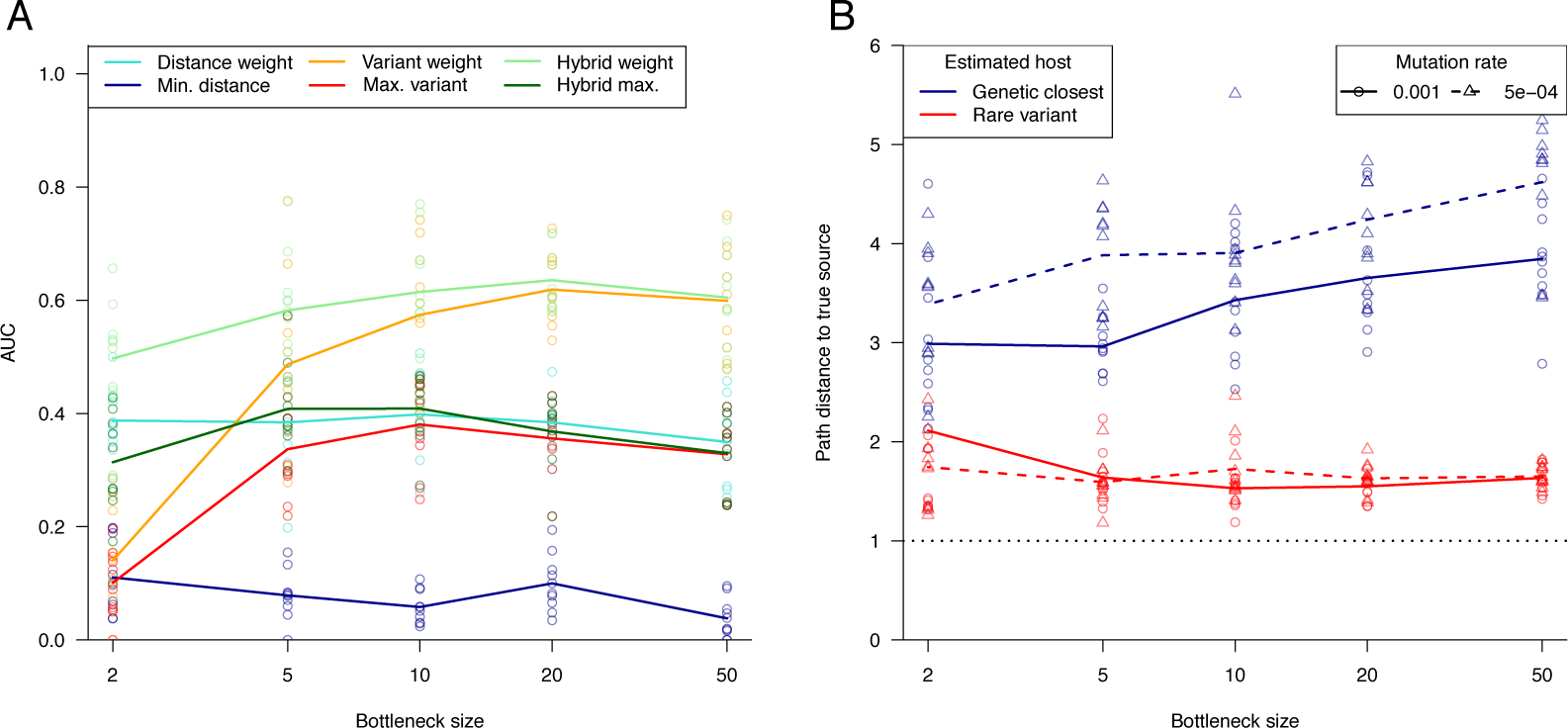
Transmission tree reconstruction accuracy for imperfectly sampled outbreaks. Data were simulated in which 30% of cases were not observed in order to assess the impact on transmission route identification. (A) The area under the ROC (AUC) metric provides an overall measure of network accuracy. Results for a mutation rate of 0.001 are shown here. (B) The true path distance between estimated transmission pairs gives insight into the extent to which transmission links are misspecified. A perfect reconstruction would have mean path length 1. Maximum variant path lengths are averaged over identified transmission pairs, that is, excluding hosts with no shared variants.

## Appendix

### Tree construction methods

Let *x*_1_,…, *x_n_* denote deep-sequence samples collected from hosts 1,…,*n*. For each sample *x_i_*, let 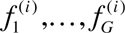 be the frequency of the majority nucleotide at loci 1,…, *G*, such that polymorphisms exist where 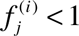. For each host, identify the set of polymorphisms 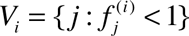. Now calculate the variant score *S_ij_* between each pair of hosts *i* and *j* to be the number of SVs belonging to the samples *x_i_* and *x_j_*; *S_ij_* = |*V_i_* ∩ *V_j_*|. If we allow for the possibility of different mutations at a given locus, we must further restrict to the set of variant positions sharing the same mutant nucleotide. The matrix (*S_ij_*)_*i*,*j*≤*n*_ can then be transformed into a weighted adjacency matrix defining an estimated transmission tree (which we call the *weighted variant tree*), in which the weight for an arrow from *i* to *j* is 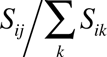. While the symmetry of *S_ij_* implies edges between a given pair of nodes must exist in both directions, these may not be of equal weight. We further define the *maximum variant tree*, in which we identify for each host the individual sharing the greatest number of variants. If multiple individuals share the maximum number of variants, these are attributed equal weight. If no individual shares any variant with a host, it is not assigned a source.

These approaches are analogous to existing methods using single genome samples. For instance, the *minimum distance tree* (or equivalently, the minimum spanning tree) is defined by assigning the source to be the individual carrying the most genetically similar sample (i.e. fewest number of SNPs). A similar approach in which temporal restrictions were additionally implemented was described in (6). Similarly, the *weighted distance tree* is defined by weighting each network edge by inverse genetic distance, such that more similar samples are given a greater weight. Variations of this method were explored in (2).

Finally we propose two hybrid approaches. In some cases, a host may share no variants with any other host in the population, such that the maximum and weighted variant approaches assign no weight to any potential sources of infection. As such, we may instead attempt to draw information from genetic distance measures where no SVs exist. The *hybrid maximum tree* and the *hybrid weighted tree* attribute sources to hosts lacking SVs according to the minimum distance and weighted distance approaches respectively. Genetic distance can be calculated using the consensus sequence of the deep sequenced sample, or with additional single genome samples. We tried both approaches, and found little difference in average performance. As such, we present results using consensus sequences only.

### Simulations

We simulated outbreaks using the R package *seedy* v1.2 (36), introducing a single infectious individual into a susceptible, homogeneously mixing population of size 100, retaining only those epidemics with final size at least 50. Genomic samples (perfectly observed deep sequence samples) were generated at a random time during each individual’s infectious period. We constructed the consensus sequence for each sample, and additionally sampled single genomes from each host in order to compare transmission route estimation using each type of data. Infection dynamics were simulated under a standard SIR (susceptible-infected-removed) model, with *R*_0_ = 2. Multiple infections were not permitted. Infections were generated by selecting *n_B_* genotypes at random from the source’s pathogen population, and allowing this inoculum to grow within the new host under neutral evolution. We varied transmission bottleneck size *n_B_* as well as mutation rate in order to simulate a range of different outbreaks. Transmission trees were visualized using the igraph package in R (37). We allowed mutation rates of 0.00025, 0.0005 and 0.001 per genome per pathogen generation, which are approximately in line with rates found for some bacterial pathogens. *Staphylococcus aureus*, for instance, has a mutation rate of 0.0005 (assuming a rate of 33×10^−6^ per nucleotide per year (38, 39) and a generation time of 30 minutes (40, 41)).

Few estimates exist for transmission bottleneck sizes, and those that do exist exhibit uncertainty – Emmett et al. estimated Ebola virus bottleneck sizes in the range of 1-800 viral particles (17), while estimates for influenza range between 100 and 250 (21). In contrast, it is thought that *Clostridium difficile* is associated with bottleneck size close to 1 (42). In our simulation studies, we explored the lower end of this range, allowing the bottleneck size to vary between 1 and 50. In practice, we found little change in outcome as the bottleneck size was allowed to be larger than this value. It is worth noting that while our simulated transmission events are a random sample of the donor’s pathogen population, in reality this may not be the case. As such, a small randomly drawn bottleneck size may be equivalent in terms of diversity to a much larger non-random sample.

### Measuring reconstruction accuracy

We used various metrics to compare the performances of the transmission route identification methods. We assumed that infection and removal times were not observed from the simulated outbreaks, and investigated the ability of the genomic data alone to contribute to transmission route identification. We used a variety of metrics to measure both the overall accuracy of the reconstructed tree, as well as the reliability of individually estimated transmission routes.

The receiver operating characteristic (ROC) curve describes the change in false positive and true positive rate for identifying a source of infection as the weight threshold for this identification varies between 0 and 1. The area under the ROC curve (AUC) is a summary statistic of this function, measuring the overall discriminatory power of the tree reconstruction, in which values closer to 1 indicate a more accurate network (43). For the unweighted reconstructions ( *minimum distance* and *maximum similarity tree*), we calculated the path distance in the true network for each proposed transmission link (ignoring directionality of edges). For instance, if we identify the route A-B, and in reality the transmission chain was A-C-B, the path distance in the true network is 2. A perfect reconstruction would thus have a mean path distance of 1. While the ROC curve treats edges as either correct or incorrect, the latter metric provides a measure of the extent to which false links are misleading (i.e. an incorrect edge with a true path length of 2 is better than an incorrect edge with a true path length of 10). This allows us to determine the risk of classifying a pair of individuals as a transmission pair, when in reality there exist intermediate hosts in the chain.

Finally, we considered the mean weight attributed to the true source of infection across an outbreak. While the path distance metric did not factor in hosts for whom no source could be attributed (due to a lack of SVs), this measure includes such hosts with a weight of zero.

### Data

We applied the SV approach to identifying potential transmission routes during the 2014 Ebola outbreak in Sierra Leone. We used samples collected from 78 patients in May-June 2014, representing a large proportion of the earliest cases in the country, sequenced to approximately 2000x coverage (19). Furthermore, we considered samples from a further 150 patients collected between June and December in the same country, sequenced with a median coverage of 374x (24). Further details of data collection, sequencing and variant calling are described in the respective original studies.

### Impact of Mutational Hotspots

To this point, simulations were performed under the assumption that mutations accumulated uniformly across the genome. However in reality polymorphisms are often concentrated in certain regions as a consequence of diversifying selection or a proportion of nucleotides which experience a heightened mutation rate. In order to explore the impact of so-called ‘mutational hotspots’, we repeated simulations with the mutation rate increased by a factor of up to 1000 in a small proportion of the genome (0.1%). As this hypermutation factor increased, a much larger proportion of samples shared at least one variant with one other case (Figure S2A), but a proportion of these SVs did not occur between true transmission pairs which impedes our ability to detect the true transmission routes, particularly when combined with small bottleneck sizes (Figure S2B). While distance-based methods were largely unaffected by the presence of hotspots, variant-based methods continued to outperform them for larger bottleneck sizes (>10) (Figure S3). In fact, for some metrics, there was a slight increase in accuracy under low bottleneck conditions with a hypermutation factor of 100 as the increased probability of mutations occurring and being transmitted outweighed the misclassification of routes due to homoplasy (Figures S4B, S4E). However, this effect was lost with a hotspot factor of 1000.

## References

1 Cottam EM, Thébaud G, Wadsworth J, et al. Integrating genetic and epidemiological data to determine transmission pathways of foot-and-mouth disease virus. Proc R Soc B 2008;275(1637):887–95.

2 Worby CJ, Lipsitch M, Hanage WP. Within-Host Bacterial Diversity Hinders Accurate Reconstruction of Transmission Networks from Genomic Distance Data. PLoS Comp Biol 2014;10(3):e1003549.

3 Didelot X, Gardy J, Colijn C. Bayesian analysis of infectious disease transmission from whole genome sequence data. Mol Biol Evol 2014;31(7): 1869–79.

4 Ypma RJF, Bataille AMA, Stegeman A, et al. Unravelling transmission trees of infectious diseases by combining genetic and epidemiological data. Proc R Soc B 2012;279:444–50.

5 Jombart T, Eggo RM, Dodd PJ, et al. Reconstructing disease outbreaks from genetic data: a graph approach. Heredity 2011;106(2): 383–90.

6 Ypma RJF, van Ballegooijen WM, Wallinga J. Relating phylogenetic trees to transmission trees of infectious disease outbreaks. Genetics 2013;195(3): 1055–62.

7 Jombart T, Cori A, Didelot X, et al. Bayesian Reconstruction of Disease Outbreaks by Combining Epidemiologic and Genomic Data. PLoS Comp Biol 2014;10(1):e1003457.

8 Streulens MJ, Deplano A, Godard C, et al. Epidemiologic typing and delineation of genetic relatedness of methicillin-resistant Staphylococcus aureus by macrorestriction analysis of genomic DNA by using pulsed-field gel electrophoresis. J Clin Microbiol 1992;30(10): 2599–605.

9 Strommenger B, Braulke C, Heuck D, et al. spa Typing of Staphylococcus aureus as a Frontline Tool in Epidemiological Typing. J Clin Microbiol 2008;46(2): 574–81.

10 Koreen L, Ramaswamy SV, Graviss EA, et al. spa Typing Method for Discriminating among Staphylococcus aureus Isolates: Implications for Use of a Single Marker To Detect Genetic Micro- and Macrovariation. J Clin Microbiol 2004;42(2): 792–9.

11 Gardy JL, Johnston JC, Ho Sui SJ, et al. Whole-Genome Sequencing and Social-Network Analysis of a Tuberculosis Outbreak. New Engl J Med 2011;364(8): 730–9.

12 Bryant JM, Schürch AC, van Deutekom H, et al. Inferring patient to patient transmission of Mycobacterium tuberculosis from whole genome sequencing data. BMC Infect Dis 2013;13:110.

13 Walker TM, Lalor MK, Broda A, et al. Assessment of Mycobacterium tuberculosis transmission in Oxfordshire, UK, 2007–12, with whole pathogen genome sequences: an observational study. The Lancet Respiratory Medicine 2014;2(4): 285–92.

14 Worby CJ, Chang H-H, Hanage WP, et al. The distribution of pairwise genetic distances: a tool for investigating disease transmission. Genetics 2014;198(4): 1395–404.

15 Hughes J, Allen RC, Baguelin M, et al. Transmission of Equine Influenza Virus during an Outbreak Is Characterized by Frequent Mixed Infections and Loose Transmission Bottlenecks PLoS Path 2012;8(12):e1003081.

16 Murcia PR, Hughes J, Battista P, et al. Evolution of an Eurasian Avian-like Influenza Virus in Naïve and Vaccinated Pigs PLoS Path 2012;8(5):e1002730.

17 Emmett KJ, Lee A, Khiabanian H, et al. High-resolution Genomic Surveillance of 2014 Ebolavirus Using Shared Subclonal Variants. PLOS Currents Outbreaks 2015(Feb 9 Edition 1).

18 Balloux F. Demographic influences on bacterial population structure. In: Robinson DA, Falush D, Feil EJ, eds. Bacterial Population Genetics in Infectious Diseases: John Wiley & Sons Inc., 2010.

19 Gire SK, Goba A, Andersen KG, et al. Genomic surveillance elucidates Ebola virus origin and transmission during the 2014 outbreak. Science 2014;345(6202): 1369–72.

20 Stack JC, Murcia PR, Grenfell BT, et al. Inferring the inter-host transmission of inflluenza A virus using patterns of intra-host genetic variation. Proc R Soc B 2013;280(1750):20122173.

21 Poon LLM, Song T, Rosenfeld R, et al. Quantifying influenza virus diversity and transmission in humans. Nat Genet 2016.

22 Paterson GK, Harrison EM, Murray GGR, et al. Capturing the cloud of diversity reveals complexity and heterogeneity of MRSA carriage, infection and transmission. Nature Communications 2015;6(6560).

23 Wertheim JO, Leigh Brown AJ, Hepler NL, et al. The global transmission network of HIV-1. J Infect Dis 2014;209(2): 304–13.

24 Park DJ, Dudas G, Wohl S, et al. Ebola Virus Epidemiology, Transmission, and Evolution during Seven Months in Sierra Leone. Cell 2015;161(7): 1516–26.

25 Pybus OG, Rambaut A. Evolutionary analysis of the dynamics of viral infectious disease. Nat Rev Genet 2009;10:540–50.

26 WHO Ebola Response Team. Ebola Virus Disease in West Africa - The First 9 Months of the Epidemic and Forward Projections. New Engl J Med 2014;371(16): 1481–95.

27 Snitkin ES, Zelazny AM, Thomas PJ, et al. Tracking a Hospital Outbreak of Carbapenem-Resistant Klebsiella pneumoniae with Whole-Genome Sequencing. Sci Transl Med 2012;4(148): 148ra16.

28 Spada E, Sagliocca L, Sourdis J, et al. Use of the minimum spanning tree model for molecular epidemiological investigation of a nosocomial outbreak of hepatitis C virus infection. J Clin Microbiol 2004;42(9): 4230–6.

29 Varble A, Albrecht RA, Backes S, et al. Influenza A virus transmission bottlenecks are defined by infection route and recipient host. Cell Host Microbe 2014;16(5): 691–700.

30 Joseph SB, Swanstrom R, Kashuba ADM, et al. Bottlenecks in HIV-1 transmission: insights from the study of founder viruses. Nat Rev Microbiol 2015;13(7): 414–25.

31 Bar KJ, Li H, Chamberland A, et al. Wide Variation in the Multiplicity of HIV-1 Infection among Injection Drug Users. J Virol 2010;84(12): 6241–7.

32 Stadler T, Kühnert D, Rasmussen DA, et al. Insights into the early epidemic spread of Ebola in Sierra Leone provided by viral sequencing. PLOS Currents Outbreaks 2014(Oct 6 Edition 1).

33 Hoenen T, Safronetz D, Groseth A, et al. Mutation rate and genotype variation of Ebola virus from Mali case sequences. Science 2015;348(6230): 117–9.

34 Rocha EP, Smith JM, Hurst LD, et al. Comparisons of dN/dS are time dependent for closely related bacterial genomes. J Theor Biol 2006;21(239(2)):226–35.

35 Leggett HC, Cornwallis CK, West SA. Mechanisms of Pathogenesis, Infective Dose and Virulence in Human Parasites. PLoS Path 2012;8(2):e1002512.

36 Worby CJ, Read TD. ‘SEEDY’ (Simulation of Evolutionary and Epidemiological Dynamics): An R Package to Follow Accumulation of Within-Host Mutation in Pathogens PLoS One 2015;10(6):e0129745.

37 Csardi G, Nepusz T. The igraph software package for complex network research. InterJournal Complex Systems 2006:1695.

38 Harris SR, Feil EJ, Holden MTG, et al. Evolution of MRSA during hospital transmission and intercontinental spread. Science 2010;327(5964): 469–74.

39 Young BC, Golubchik T, Batty EM, et al. Evolutionary dynamics of Staphylococcus aureus during progression from carriage to disease. Proc Natl Acad Sci USA 2012;109:4550–5.

40 Chang-Li X, Hou-Kuhan T, Zhau-Hua S, et al. Microcalorimetric study of bacterial growth. Thermochimica Acta 1988;123:33–41.

41 Dengremont E, Membré JM. Statistical Approach for Comparison of the Growth Rates of Five Strains of Staphylococcus aureus. Appl Environ Microbiol 1995;61(12): 4389–95.

42 Didelot X, Eyre DW, Cule M, et al. Microevolutionary analysis of Clostridium difficile genomes to investigate transmission. Genome Biology 2012;13(12):R118.

43 Fawcett T. An introduction to ROC analysis. Pattern Recog Lett 2006;27(8): 861–74.

